# Targeting Telomere Integrity Enhances Radiosensitivity and Reveals Adaptive DNA Damage Response Vulnerabilities in Diffuse Midline Glioma

**DOI:** 10.64898/2026.11.13.766496

**Authors:** Banlanjo Umaru, Zhuang Wang, Deepak Kumar Mishra, Shiva Senthil Kumar, Archana Mahadevan Iyer, Komal Khan, Hao-Han Pang, Sergei Gryaznov, Maryam Fouladi, Rachid Drissi

## Abstract

Diffuse midline glioma (DMG), including diffuse intrinsic pontine glioma (DIPG), remains one of the most lethal pediatric malignancies, with limited durable benefit from radiotherapy (RT) due in part to adaptive DNA damage response (DDR) mechanisms. Here, we investigated whether targeting telomere integrity with the telomerase substrate analogue 6-thio-2′-deoxyguanosine (6-thio-dG; THIO) could induce therapeutically exploitable DNA damage in patient-derived DIPG models. THIO selectively suppressed DIPG cell proliferation while sparing telomerase-negative normal diploid fibroblasts and induced telomere dysfunction, as evidenced by increased telomere dysfunction-induced foci (TIFs), γH2AX accumulation, G_2_/M checkpoint arrest, and apoptosis. In orthotopic DIPG xenografts, THIO increased intratumoral TIF formation, suppressed tumor growth, and prolonged survival. Transcriptomic profiling further revealed activation of cellular stress-response pathways and downregulation of *E2F* genes, mitotic, and metabolism-associated gene programs. Mechanistically, THIO-induced progressive genomic and telomeric damage was associated with sequential activation of ATM and ATR signaling, consistent with engagement of adaptive DDR pathways. Combining THIO with ionizing radiation (IR) produced synergistic antitumor effects across three independent DIPG models representing distinct H3 mutation subtypes, with increased G_2_/M accumulation, genomic and telomeric DNA damage, and apoptotic cell death. *In vivo*, THIO combined with IR significantly improved therapeutic efficacy and prolonged median survival compared with monotherapy, including RT alone. Notably, pharmacologic inhibition of ATM or ATR attenuated THIO-induced checkpoint arrest and increased apoptosis, implicating DDR checkpoint signaling as an adaptive survival dependency following telomere damage. Collectively, these findings identify telomere-directed DNA damage as a promising therapeutic approach for DMG and provide a mechanistic rationale for combining THIO with RT and DDR checkpoint inhibition.

## INTRODUCTION

Diffuse midline gliomas (DMGs), including diffuse intrinsic pontine glioma (DIPG), are a deadly class of pediatric high-grade gliomas (pHGGs) that account for 10%-15% of all pediatric brain tumors^1–3^. DIPG is associated with a dismal prognosis, with a median overall survival (OS) of 9-11 months and a 2-year survival of less than 10%^4^. Its diffusely infiltrative growth pattern and location within the brainstem, where critical neurological structures are concentrated, preclude safe surgical resection^5,6^. Fractionated radiotherapy (RT) remains the standard of care, providing transient clinical benefit and extending OS by only 3-4 months on average^7,8^. Despite multimodal therapies, including the recently approved dordaviprone for H3K27-mutant DMG^9^, durable disease control remains elusive, and survival outcomes have not changed substantially over the past several decades^10–12^. These challenges underscore the urgent need for more effective therapeutic approaches, including rationally designed combination strategies to improve treatment response and prolong survival.

A major barrier to durable treatment response in DMG, and a key contributor to radio-resistance, is the capacity of tumor cells to withstand endogenous and therapy-induced DNA damage through activation of the DNA damage response (DDR). This response engages ATM- and ATR-dependent signaling, cell-cycle checkpoints, and DNA repair pathways that collectively promote tumor cell survival despite genotoxic stress^13,14^. Telomeres represent a particularly vulnerable chromosomal substrate at which persistent DNA damage can accumulate, ultimately triggering tumor cell death. Telomeres are composed of several kilobases of tandem TTAGGG repeats and play an essential role in maintaining chromosomal stability by physically capping the ends of eukaryotic linear chromosomes^15^. We and others have demonstrated that replicative senescence-associated telomere shortening induces telomere dysfunction, thereby activating ATM-dependent DDR signaling^16,17^. This response is characterized by the formation of telomere dysfunction-induced foci (TIFs), in which DDR mediators, including phosphorylated ATM, γH2AX and 53BP1, accumulate at dysfunctional telomeres^18^. Telomere length is maintained by a telomerase, which is reactivated in approximatively 73%-90% of human cancers but remains largely inactive in most normal somatic tissues^19,20^. We have previously shown that *hTERT*, the catalytic subunit of telomerase, is expressed in over 73% of DIPG patient specimens^21^. Furthermore, our recent findings demonstrated that patients with pHGG harboring telomerase-positive tumors have significantly shorter event-free survival and overall survival than those with telomerase-negative tumors^22^. The preferential activation of telomerase in tumors relative to most normal somatic tissues, together with its association with poor patient outcomes, positions the telomerase-telomere axis as a promising therapeutic target. We previously demonstrated that imetelstat (IMT), a potent direct inhibitor of telomerase^23,24^ had limited therapeutic efficacy as a radiosensitizer in medulloblastoma with minimal effects on tumor growth, recurrence, and survival despite effective intratumoral inhibition of telomerase activity^25^. Furthermore, our preclinical studies and a subsequent phase II clinical trial evaluating IMT in children with recurrent central nervous system malignancies demonstrated limited clinical efficacy and poor tolerability, in part due to treatment-related toxicities that precluded more frequent dosing^26^. Collectively, these findings highlight the limitations of direct telomerase inhibition as a therapeutic strategy and underscore the need for alternative telomere-targeting approaches capable of inducing rapid and sustained telomere dysfunction.

6-thio-2′-deoxyguanosine (6-thio-dG; THIO) represents a mechanistically distinct therapeutic strategy approach that exploits telomerase activity to selectively induce telomere dysfunction in cancer cells. Following intracellular metabolism to 6-thio-dGTP, THIO recognized by telomerase and incorporated into newly synthesized telomeric DNA. In a foundational study, Mender *et al.* demonstrated that the telomerase-dependent incorporation triggers rapid telomere dysfunction and DNA damage, resulting in preferential killing of telomerase-positive cancer cells while largely sparing telomerase-deficient normal cells^27^. Importantly, the therapeutic effects of THIO extend beyond direct tumor cell cytotoxicity. We have demonstrated that, in therapy-resistant pediatric brain tumor models, telomerase-mediated THIO incorporation induces TIFs, sequential activation of ATR and ATM signaling, G_2_/M cell-cycle arrest, and robust activation of DNA damage repair pathways, ultimately producing antitumor activity in high-risk group 3 medulloblastoma. Importantly, we further demonstrated that THIO crosses the blood–brain barrier in an orthotopic model of DIPG, supporting its potential as a therapeutic agent for primary brain tumors^28^. Furthermore, studies in immunocompetent tumor models have shown that THIO-induced telomere stress activates STING-dependent innate and adaptive antitumor immune responses, thereby enhancing tumor sensitivity to immune checkpoint blockade ^29–31^. Collectively, these findings establish telomere injury as more than a direct cytotoxic mechanism, supporting its potential role as a therapeutic priming event that reshapes tumor cell responses to subsequent genotoxic stress.

Here, using molecularly diverse patient-derived DIPG models and orthotopic xenografts, we investigated the cellular and therapeutic consequences of THIO-induced telomere damage in combination of radiotherapy. We demonstrated that THIO induces progressive telomere dysfunction, activates ATM/ATR checkpoint signaling, and broadly suppresses transcriptional programs associated with cellular proliferation and metabolism. Furthermore, THIO enhances radiation response *in vitro* and confers a significant survival benefit in an orthotopic DIPG model. Finally, pharmacologic inhibition of ATM and ATR reveals a functional contribution of these checkpoint pathways to the cellular response to THIO-induced telomere damage. These findings establish telomere-directed damage as a mechanistically rational strategy for enhancing RT response and uncovering DDR dependencies in DIPG.

## MATERIALS AND METHODS

### Cell lines, culture conditions, and reagents

Primary normal human foreskin fibroblasts (HFF) (ATCC CRL-2091) were purchased from the American Type Culture Collection (ATCC). HFF cells were cultured in EMEM (Gibco) supplemented with 10% FBS. Primary patient-derived DIPG cell lines CCHMC-DIPG-1 (H3-wild type), CCHMC-DIPG-2 (H3.3-K27M) and CCHMC-DIPG-IV (H3.3-K27M) were generated in the Drissi laboratory as previously described^32,33^. The DIPG cell lines SU-DIPG-XXXVI (H3.1K27M), SU-DIPG-IV (H3.1K27M) and SU-DIPG-XIII (H3.3-K27M) were generously provided by Dr. Michelle Monje (Stanford University). PBT-29FH generated from treatment-naïve patient-derived biopsy, was obtained from Dr. James Olson (Seattle Children’s Hospital). The intrauterine electroporation (IUE)-derived murine DIPG cell lines IUE-24-C5 (H3-wild type), IUE-24-B1 (H3.3-K27M), and IUE-26-D6 (H3.1K27M), were a kind gift from Dr. Timothy N. Phoenix (Cincinnati Children’s Hospital Medical Center, Cincinnati). The DIPG cells were cultured in neurosphere stem cell media according to lab established protocols as previously described^32,33^. The IUE murine cell lines were cultured in neurosphere stem cell media with addition of N2-supplement (Gibco) following previous protocols^34,35^. Using the Universal Mycoplasma Detection Kit (30-1012K; ATCC, Manassas, VA, USA), all cell lines used in this study were confirmed to be negative for mycoplasma contamination.

THIO was provided by MAIA Biotechnology, Inc. (Chicago, IL) and was dissolved in DMSO:water (1:1) to prepare a 10 mmol/L stock solution for *in vitro* studies. For *in vivo* studies, THIO was prepared in a 5% DMSO solution. Kinase inhibitors of ATM (KU-55933) and ATR (VE-822) were purchased from Selleckchem and reconstituted in DMSO.

### Telomerase activity assay

Telomerase enzyme activity was assessed by Telomeric Repeat Amplification Protocol (TRAP) assay using the TRAPeze Telomerase Detection Kit (cat# S7710, Millipore, Burlington, MA, USA). Protein extraction and TRAP assay were performed as previously described^25^. A total of 50 ng of total protein was used to assess the telomerase activity by performing polyacrylamide gel (12.5%) electrophoresis. HeLa cell line (30 ng total protein) and CHAPS buffer were used as positive and negative controls, respectively.

### Cell proliferation assay

Cell proliferation was measured using WST-1 assay (cat# MK4000, Takara Bio Inc., Shiga, Japan) as previously described^36,37^ and following the manufacturer’s instructions. WST-1 reagent was added to each well at a final dilution of 1:10, incubated for 1-2 hrs. at 37 °C, and absorbance was measured at 450 nm with 650 nm as the reference wavelength.

### Cell cycle analysis

The cell cycle analysis was performed as previously described^28^. Briefly, following treatment at indicated time point, cells were collected as single cells and fixed with ice-cold 70% ethanol, and stored at −20°C for at least 1 hr. Cells were then washed twice with PBS and stained with propidium iodide (PI) solution containing 25 µg/mL PI (Cat# P4864, Sigma Aldrich, St. Louis, MD, USA), and 100 μg/mL Ribonuclease-A (RNAase A) (Cat# 12091021, Invitrogen, Carlsbad, CA, USA) for at least 30 mins in the dark at room temperature. Flow cytometry was performed using the BD LSR Fortessa (BD Biosciences), and the data were analyzed using FlowJo v.10 software (FlowJo, Ashland, OR USA).

### Immunofluorescence (IF)

Immunofluorescence staining was performed as previously reported^32^. Briefly, treated cells were fixed with 4% paraformaldehyde (PFA), washed with PBS and permeabilized with 0.5% Triton-X-100 in PBS. Next, cells were blocked for 1 hr in 5% normal donkey serum (NDS) with 0.3% Triton X-100 in TBS. Cells were incubated overnight at 4°C with primary antibodies against γH2A.X (cat# 9718S, Cell Signaling Technology, Danvers, MA, USA) and cleaved caspase-3 (Cell Signaling Technology). Next day, cells were washed with PBS buffer and stained with secondary antibodies; Alexa Fluor 568 donkey anti-rabbit IgG (H + L) (1: 500, Cat# 711-575-152, Jackson ImmunoResearch) and Alexa Fluor 647 donkey anti-mouse IgG (H + L) (1 : 500, Cat# 715-605-151, Jackson ImmunoResearch, West Grove, PA, USA). The samples were then mounted with antifade mounting medium containing DAPI (H-1200-10, Vector Laboratories, Plain City, OH, USA) and imaged using the Nikon Ti2 Confocal Microscope (Nikon Instruments Inc.). Image processing was performed using NIS-Elements AR (v5.42.03).

### Telomere IF-FISH

For formalin-fixed paraffin-embedded (FFPE) mouse brain tissue, sections were deparaffinized in xylene, rehydrated through a graded ethanol concentration series, and subjected to heat-induced antigen retrieval for 20 mins in 10 mM Citrate buffer (pH 6.0). The tissue sections were then dehydrated with a graded ethanol concentration series and air dried. Cultured cells were fixed and permeabilized with 0.5% Triton-X-100 in PBS. Next, both the tissue sections and cells were covered with hybridization solution containing 70% formamide, 0.5% blocking reagent (Roche Diagnostics) diluted in 100 mmol/L maleic acid and 150 mmol/L sodium chloride, and 10 mmol/L Tris (pH 7.5), and 100 ng/mL TelC-Cy3 PNA telomere probe (Biosynthesis). Samples were denatured for 6 mins at 84°C and hybridized for at least 2 hrs. at room temperature. After washing three times with 70% formamide and 10 mmol/L Tris (pH 7.5) and then three times with TBS, samples were blocked in 5% NDS and 0.3% Triton X-100 in TBS for 1 hr and incubated overnight at 4 °C with primary antibodies against γH2A.X (cat# 9718S, Cell Signaling Technology) for cells or 53BP1 (NOVUS NB100-304) for tissue sections. Next day, they were incubated with secondary antibody Alexa Fluor 488 donkey anti-rabbit IgG (H + L) (1 : 500, Cat# 711-545-152, Jackson ImmunoResearch). Finally, the samples were mounted with DAPI-containing mounting medium and imaged using a Nikon Ti2 confocal microscope. Image processing was done with NIS-Elements AR (v5.42.03).

### Western blotting

Western blotting was performed as previously described^33^. Antibodies used were against ATM-S1981-P (cat# 200-301-400, Rockland), ATM (cat# A1106, Sigma), ATR-T1989-P (cat# GTX128145, GeneTex, Irvine, CA, USA), ATR (cat# 13934S), CHK2 (cat# 2662S), CHK2-Thr68-P (cat# 2661S), CHK1 (cat# 2360S), CHK1-S345-P (cat# 2348S), γH2A.X (cat# 9718S), β-Actin (cat# 3700S), and Histone H3 (cat# 3638S) from Cell Signaling Technology (Danvers, MA, USA). Bands visualized with ECL substrates and were captured using the ChemiDoc MP imaging system (Bio-Rad Laboratories, Hercules, CA, USA) and quantified using ImageJ software (National Institutes of Health, Bethesda, MD, USA).

### *hTERT* Expression by qRT-PCR

qRT-PCR analysis was performed as previously described^21^. Briefly, total RNA extraction was performed from cell lysates using the Qiagen RNeasy Mini Kit (Qiagen) and cDNA was synthesized using the High-Capacity RNA-to-cDNA Kit (Applied Biosystems). Primer/probe sets for *hTERT* (Hs00972650_m1), *TBP* (Hs00427621_m1) and *GAPDH* (Hs03929097_g1) were purchased from Applied Biosystems. Two independent qRT-PCR experiments were performed with each sample run in triplicates. Changes in gene expression (expressed as RQ) were calculated using the ΔΔCt method relative to HFF.

### RNA sequencing

RNA sequencing procedure was performed as previously described^38^. RNA extraction was performed using RNeasy Plus Mini Kit (Cat# 74134, Qiagen, Hilden, Germany) following the manufacturer’s instructions and as previously described^39^. RNA concentration was quantified using Invitrogen Qubit 4 Fluorometer and RNA BR assay kit (Cat# Q10211, Invitrogen). Transcript abundances were quantified using Salmon (v1.11.0) with the GRCh38 reference transcriptome (GENCODE release 21). Downstream analysis was performed on R (v4.5) after importing quantification files. Briefly, the DESeq2 pipeline (v1.52.0)^40^ was used for differential expression analysis. THIO versus vehicle comparisons were performed for each cell line using the Wald test with all the samples fitted into a single model. Gene expression data were normalized as TPM values and subjected to a log₂(TPM + 1) transformation for downstream visualization. Pathway enrichment was evaluated using clusterProfiler (v4.2.2) and GSEA with gene sets from the Molecular Signature Database (MSigDB, v2024.1.Hs). The p-values were adjuste^40^d for multiple testing using the Benjamini-Hochberg method.

### Synergy analysis

Synergy analyses were performed following cell proliferation analysis by WST-1 assay as previously indicated. Patient-derived DIPG cells were plated in 96-well plates at a density of 2000 - 5000 cells per well depending on cell-specific kinetics. For THIO and ionizing radiation (IR) co-treatments, cells were treated with sequentially diluted concentrations of THIO (DMSO, 0.01, 0.1, 0.5, and 10 µM) and incubated at 37 °C for 48 hrs., then subjected to different doses of X-ray (0, 1, 2, 4, 8, and 10 Gy) using the using X-Rad320 (Precision X-Ray, Madison, CT, USA). The absorbance was measured 72 hrs. after incubation. For THIO and ATM inhibitor (ATMi) or ATR inhibitor (ATRi) co-treatments, 24 hrs. after plating, cells were treated with either ATMi (0.5, 1, 10, 20, and 50 µM) or ATRi (10, 20, 50, 100, and 500 nM) and incubated at 37 °C for 2 hrs., and then with (DMSO, 0.01, 0.1, 0.5, and 10 µM). The absorbance was measured 72 hrs. after incubation at 37 °C. The dose-response curves were generated using GraphPad Prism 10. The synergistic effect was analyzed using SynergyFinder 3.0^41^. A peak bliss synergy scores greater than 10 was indicative of a synergistic effect. All assays were performed with three independent biological replicates.

### Immunohistochemistry

Formalin-fixed paraffin-embedded (FFPE) mouse brain tissue sections were deparaffinized in xylene followed by rehydration through a graded ethanol concentration series and subjected to heat-induced antigen retrieval for 20 mins in 10 mM citrate buffer (pH 6.0). Endogenous peroxidase activity was quenched with 1% H_2_O_2_ followed by washing, then blocking with 10% goat serum in TBST for 1 hr and incubated overnight at 4 °C with primary antibody cleaved caspase-3 (1:1,000; rabbit; Cell Signaling Technology) in 2% goat serum. After washing 3x with TBST, slides were treated with biotinylated anti-rabbit secondary antibody (1:500), amplified using the ABC Kit (Vector Laboratories). The signal was visualized with DAB (Vector Laboratories), and Harris Hematoxylin (Sigma) counterstained. H&E staining was performed with hematoxylin-1 and eosin-Y (Thermo Fisher Scientific). Slides were mounted with Permount (Thermo Fisher Scientific) and imaged with the Nikon Eclipse Ti2-E-Widefield Microscope (Nikon Instruments Inc.). IHC stained slides were quantified using Image J.

### Orthotopic xenograft studies and bioluminescence imaging

Athymic Ncr-*nu/nu* female mice (J:NU (*Foxn1^nu^/Foxn1^nu^*) were purchased from the Jackson Laboratory and were maintained in the Animal Resource Core at Nationwide Children’s Hospital. All animal studies were conducted under protocols approved by the Institutional Animal Care and Use Committee (IACUC) at Nationwide Children’s Hospital and Ohio State University, and in accordance with the NIH guidelines for the care and use of laboratory animals. The number of mice in each treatment group is indicated in the figure legends. Four to five weeks old mice were intracranially implanted with luciferase-tagged PBT-29FH (200,000 cells) or SU-DIPG-XXXVI (100,000 cells) in 4 μL PBS using a stereotactic surgical setup with coordinates 0.8 mm lateral and 1 mm posterior to lambda and 4 mm deep, corresponding to the pons location in mouse brain. Mice were randomized to treatment groups 7-10 days post implantation based on bioluminescence imaging (BLI). THIO was administered by intraperitoneal (i.p.) injection following the indicated schemes and description in the figure legends. For BLI, mice were anesthetized with isoflurane (Patterson Veterinary Supply, #07-890-8115), i.p injected with D-luciferin (Revvity, #122799-5), and imaged using the IVIS Spectrum Imaging System (Perkin Elmer). Weekly BLI was performed to measure the tumor signals. Radiance analysis was performed using the Living Image Software (Revvity).

### Radiation delivery

*In vitro* irradiation studies were performed using the X-RAD 320 irradiator (Precision X-Ray, Inc.), at Preclinical Behavior & Imaging Core in Nationwide Children’s Hospital, with a source-to-surface distance of 60 cm and a dose rate of ∼2.0 Gy/min. Cells were irradiated 48 hrs. post THIO treatment with different doses of X-ray (0, 1, 2, 4, 8, and 10 Gy) depending on the experimental set up as described in the results. For *in vivo* animal irradiation, the small animal radiation research platform (SARRP) at the Small Animal Imaging Core at The Ohio State University was used to deliver fractionated radiation to pons area of the brain. Irradiation was image-guided using computerized tomography (CT) imaging and a rotating X-ray source with adjustable collimator to deliver specific radiation dose and minimize exposure to non-targeted tissues and organs. Three-dimensional reconstruction and MuriPlan software were used for treatment planning. The scheme in Figure 4 indicates delivery of 10 Gy per radiation cycle in 2 Gy daily fractions for a total 3 radiation cycles (30 Gy cumulative).

### Statistical analysis

The statistical analysis of RNA sequencing data was performed using the intrinsic methods implemented within the respective R packages. Downstream analysis, including differential gene expression and pathway enrichment, was carried out using default statistical frameworks embedded in each package. Statistical analyses for other experimental assays were performed in GraphPad Prism v10 unless otherwise stated. Two group comparisons were analyzed using unpaired two-tailed Student’s t-test, and comparisons involving more than two groups were analyzed by one-way or two-way ANOVA followed by appropriate post hoc tests. Experiments were repeated at least twice. Survival data were analyzed using the Mantel-Cox log-rank test, by first running a global log-rank test on whether the survival curves were different from each other and then comparing individual treatment arms post hoc using the log-rank test. Data are presented as mean±standard error of the mean (SEM). P-values from the t-test, multiple comparison test, and log-rank test are reported as follows: ns, not significant, *P < 0.05, **P < 0.01, ***P < 0.001, and ****P < 0.0001.

## RESULTS

### THIO suppresses DIPG cell growth across molecular subtypes and induces progressive telomere dysfunction

We first investigated the response of a molecularly diverse panel of patient-derived DIPG cells to THIO. The panel comprised telomerase-positive H3-wild-type cell line: CCHMC-DIPG-1, H3.3K27M cell lines: CCHMC-DIPG-2, CCHMC-DIPG-IV, SU-DIPG-XIII and PBT-29FH, and H3.1K27M cell lines: SU-DIPG-IV and SU-DIPG-XXXVI (Figure S1A-B). THIO induced concentration-dependent inhibition of proliferation across all patient-derived DIPG models tested. In contrast, telomerase-negative normal human diploid fibroblasts (HFFs), exhibited substantially greater resistance with an IC_50_ of 14.8 µM, exceeding the highest concentration tested (Figure 1A). Comparable sensitivity was also observed across genetically defined murine H3-wild-type, H3.3K27M, and H3.1K27M DIPG models (Figure S1C-D), indicating that THIO-mediated growth inhibition was not restricted to a specific H3 variant or K27M mutation subtype.

**Figure 1:**
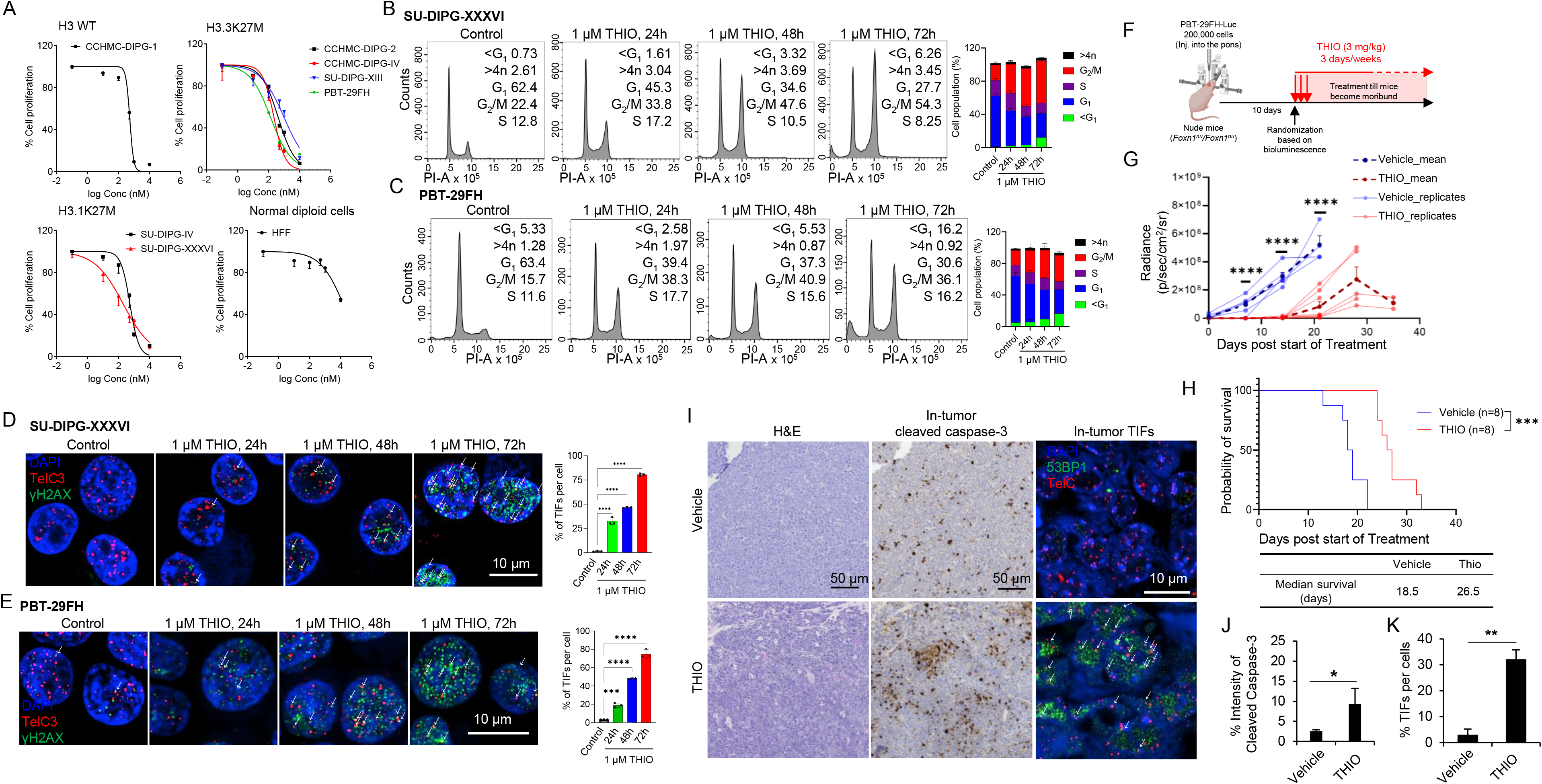
THIO induces telomere dysfunction and inhibits DIPG growth *in vitro* and *in vivo*. **(A)** Dose-response curves showing the effects of THIO on H3-wild-type, H3.3K27M, and H3.1K27M patient-derived DIPG cells and normal diploid human foreskin fibroblasts (HFFs). Cell proliferation was assessed 72 hrs following THIO treatment. **(B–C)** Flow cytometric analysis of cell cycle distribution in SU-DIPG-XXXVI and PBT-29FH cells following treatment with 1 μM THIO for 24–72 hrs. Quantification of cell cycle phase distribution is shown to the right of the representative flow cytometry plots. **(D–E)** Representative telomere immunofluorescence–fluorescence in situ hybridization (IF-FISH) images showing DAPI, TelC, and γH2AX signals following THIO treatment. Arrows indicate representative telomere dysfunction-induced foci (TIFs). Graphs show quantification of TIFs per cell, with 200-500 cells analyzed per treatment condition. Error bars represent the standard deviation (SD) of at least three fields per treatment condition, with more than 100 cells analyzed per field. **(F)** Schematic representation of the experimental design for orthotopic PBT-29FH-Luc tumor implantation and subsequent THIO treatment. **(G)** Longitudinal monitoring of tumor burden by bioluminescence imaging using the IVIS system. **(H)** Kaplan–Meier survival analysis of mice bearing orthotopic PBT-29FH-Luc tumors treated with vehicle or THIO. Median survival was 18.5 days in the vehicle group and 26.5 days in the THIO-treated group. **(I)** Representative images of H&E, cleaved caspase-3, and telomere/53BP1 staining of orthotopic tumor sections. Telomeres were detected using TelC (red), 53BP1 is shown in green, and nuclei were counterstained with DAPI (blue). Arrows indicate representative TIFs. **(J-K)** Quantification of intratumoral cleaved caspase-3 and TIFs formation. Cleaved caspase-3 staining intensity was quantified using imagej and normalized to tumor area. TIFs quantification was performed across at least three fields per mouse, with more than 50 cells analyzed per field (n=3 mice). Error bars represent the standard deviation (SD). *In vitro* experiments were performed in triplicates (n=3). Statistical significance is indicated as follows: *P < 0.05, **P < 0.01, ***P < 0.001, and ****P < 0.0001.

Because dysfunctional telomeres are recognized as persistent DNA damage lesions that can impede cell cycle progression, we next investigated whether THIO treatment altered cell cycle distribution in DIPG cells. We previously demonstrated that THIO treatment induces G_2_/M cell cycle arrest in telomerase-positive HFF+ hTERT cells (HFF cells ectopically expressing *hTERT*), HeLa cells, and MB004 primary high-risk group 3 medulloblastoma cells^28^. Similarly, treatment of patient-derived DIPG cells with 1 μM THIO resulted in a significant, time-dependent accumulation of cells in the G_2_/M phase, consistent with cell cycle arrest (Figure 1B-C and S1E). Notably, prolonged THIO exposure (72 hrs.) also led to an increase in the sub-G_1_ population, consistent with the transition from cell cycle arrest toward cell death (Figure 1B-C, S1E). We next directly assessed THIO-induced telomere dysfunction-induced foci (TIFs). Immunofluorescence-fluorescence *in situ* hybridization (IF-FISH) staining for telomeres and the DDR marker γH2AX revealed that treatment with 1 μM THIO induced a significant, time-dependent increase in DNA damage and TIFs formation (Figure 1D-E, S1F). TIFs were detectable as early as 24 hrs. after treatment and increased markedly at 48 and 72 hrs. (Figure 1D-E, S1F). Together, these findings demonstrate that molecularly diverse patient-derived DIPG models are sensitive to THIO, with the cellular response characterized by early onset of telomere dysfunction, and by progressive cell cycle checkpoint activation and accumulation in the sub-G_1_ population, consistent with subsequent cell death.

### THIO induces anti-tumor activity and telomere dysfunction in orthotopic DIPG models

To determine whether the sensitivity of patient-derived DIPG models to THIO observed *in vitro* could be translated to an orthotopic *in vivo* setting, luciferase-labeled PBT-29FH cells were stereotactically implanted into the pons of immune deficient athymic mice. Following confirmation of tumor formation by bioluminescence imaging (BLI), animals were randomized to receive either vehicle or THIO (3 mg/kg) administered once daily for three consecutive days each week until endpoint criteria were met (Fig. 1F). THIO treatment resulted in significant suppression of tumor growth compared with vehicle treatment (Figure 1G). This suppression of tumor growth translated into a significant survival benefit, with a median survival increasing from 18.5 days in vehicle-treated mice to 26.5 days in THIO-treated mice following treatment initiation (log-rank P=0.0002; Figure 1H). Immunohistochemistry staining provided pharmacodynamic evidence of THIO activity within intracranial tumors. THIO-treated tumors exhibited a significant increase in cleaved caspase-3 staining, consistent with enhanced apoptosis, as well as a significant increase in 53BP1/telomere colocalization, indicative of TIFs (Figure 1I-K). Collectively, these findings demonstrate that systemically administered THIO suppresses orthotopic DIPG tumor growth, prolongs survival, and induces telomere dysfunction and apoptosis within intracranial tumors, supporting its ability to cross the blood-brain barrier and extend previous evidence of its antitumor activity against telomerase-positive pediatric brain tumors *in vivo* ^28^.

### THIO induces a coordinated cellular stress response while suppressing DNA replication, mitotic, and metabolic programs

To define the cellular programs engaged following THIO-induced telomere damage, we performed transcriptomic profiling of patient-derived DIPG cell lines, SU-DIPG-XXXVI and CCHMC-DIPG-1. Cells were treated with 0.5 µM of THIO for 48 hrs. prior to bulk RNA sequencing analysis. The principal component analysis demonstrated strong model-specific separation, with reproducible treatment-associated transcriptional shifts within each model (Figure S2A). Differential-expression analyses showed widespread remodeling of genes involved in DNA replication, chromosome segregation, and DNA damage regulation (Figure 2A-B). Cross model comparison also identified a shared core of concordantly regulated genes within the DNA replication and DNA damage regulation (Fig. S2B). Furthermore, hallmark gene-set analysis revealed a stress phenotype in THIO-treated patient-derived DIPG cells. In SU-DIPG-XXXVI, THIO positively enriched the p53 pathway, TNFα signaling through NF-κB, unfolded-protein response, and UV-response-up signatures. In parallel, programs linked to oxidative phosphorylation, glycolysis, MYC targets, fatty-acid metabolism, E2F target genes, G_2_/M checkpoint, cholesterol homeostasis, and the mitotic spindle were negatively enriched (Figure 2C). Similarly, CCHMC-DIPG-1 showed a closely related response, including activation of p53, unfolded-protein and stress-associated programs and suppression of *E2F* genes, G_2_/M, mitotic-spindle, and metabolic pathways (Figure 2D). The concurrent activation of stress-response pathways and suppression of *E2F* genes, mitotic, and metabolic programs suggests that THIO induces a transition from a proliferative state to a damage-adapted cellular state. We therefore investigated the effects of THIO on DDR signaling at the molecular level. Western blot analysis demonstrated that THIO induced time-dependent ATM activation, as evidenced by increased phosphorylation at Ser1981, across CCHMC-DIPG-1, CCHMC-DIPG-2, and SU-DIPG-XXXVI cells, accompanied by a concomitant increase in downstream CHK2 phosphorylation at Thr68. ATM and CHK2 phosphorylation progressively increased at later time points, while γH2AX accumulation was observed across all three cell models. ATR phosphorylation at Thr1989 exhibited a dynamic, time-dependent pattern across all three DIPG cell lines, reaching a peak at 24 hrs followed by a gradual decline at 48 and 72 hrs. These findings corroborate our previous observation of sequential THIO-mediated activation of ATM and ATR pathways^28^, accompanied by increased accumulation of γH2AX, a marker of DNA damage. Taken together, the transcriptomic and protein-level data reveal a conserved cellular response in which THIO-induced telomere disruption activates DNA damage signaling while concurrently suppressing transcriptional programs essential for DNA replication and mitotic progression. This response is consistent with the broader replication-stress vulnerability increasingly recognized in DMG^42,43^.

**Figure 2:**
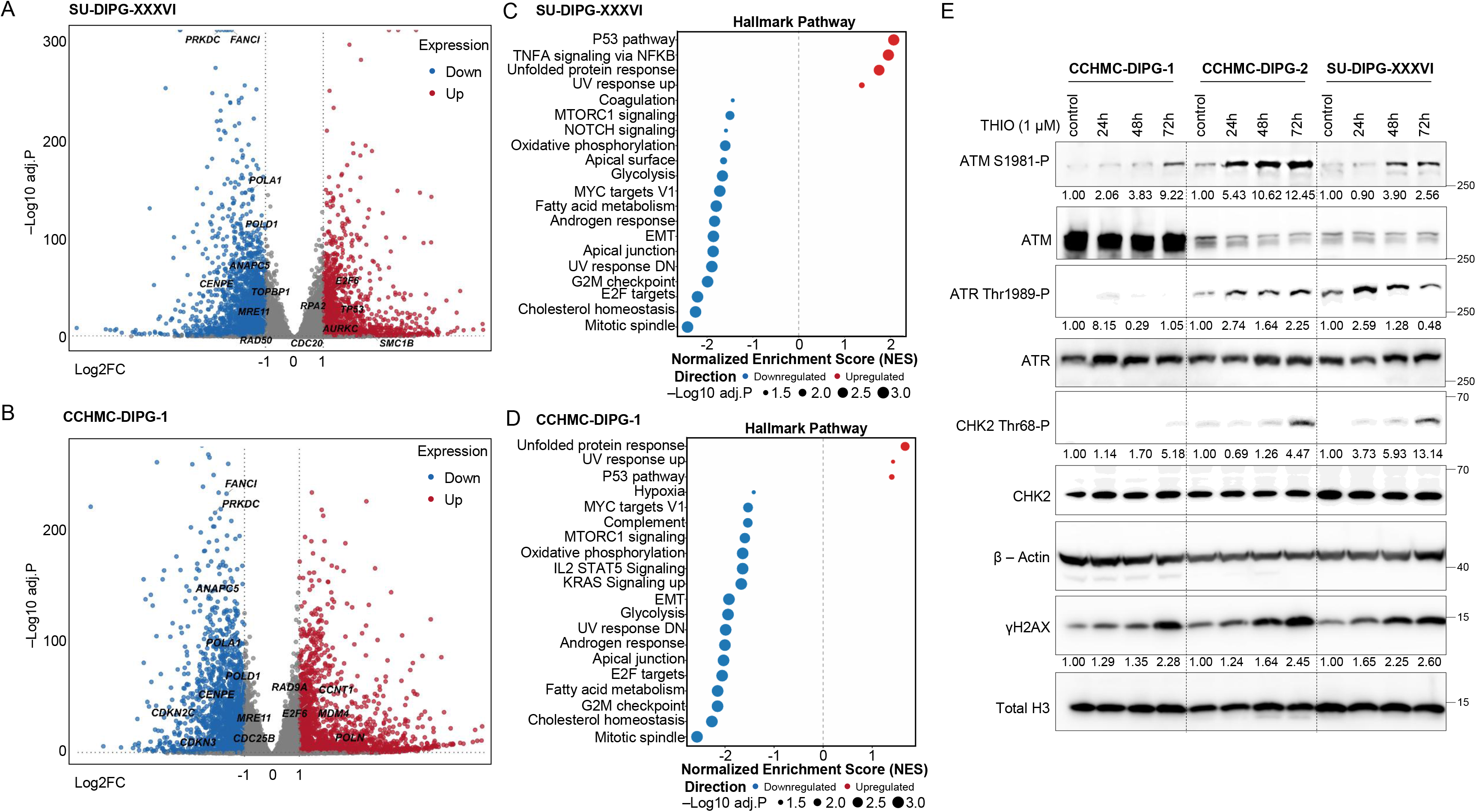
THIO induces a stress-associated transcriptional response and activates DNA damage checkpoint signaling. (**A-B)** Volcano plots depicting differentially expressed genes following THIO treatment in SU-DIPG-XXXVI and CCHMC-DIPG-1 cells. **(C-D)** Hallmark pathway enrichment analysis of transcriptional changes induced by THIO treatment in SU-DIPG-XXXVI and CCHMC-DIPG-1 cells. **(E)** Immunoblot analysis of ATM, ATR, CHK2, and γH2AX in cells treated with 1 μM THIO for 24-72 hrs. Densitometric values beneath the blots indicate relative protein expression. Phosphorylation levels of ATM, ATR, and CHK2 were normalized to their respective total protein levels and subsequently to β-actin. γH2AX levels were normalized to total histone H3.

### THIO potentiates radiation-induced DNA damage and apoptotic cell death

Given that fractionated RT remains the standard of care for DIPG, we next investigated whether THIO-induced telomere damage could potentiate radiation responses in patient-derived DIPG cell lines. To this end, we exposed three patient-derived DIPG cell lines to combination matrices of THIO and IR. Because THIO efficacy is replication-dependent and requires progression through S phase for incorporation into telomeres, administering THIO before IR was expected to maximize therapeutic efficacy. Consistent with this rationale, sequential administration of THIO prior to IR resulted in greater suppression of cell viability and proliferation than concurrent treatment (data not shown). Based on these findings, THIO was administered 48 hrs. prior to IR exposure for subsequent combination studies. The interactive effects of THIO and IR were assessed using Bliss independence scoring analysis implemented in the SynergyFinder 3.0 platform. Using the established Bliss synergy threshold of greater than 10, we identified regions of positive interaction in all three DIPG cell lines, with peak Bliss scores of 21.54 in SU-DIPG-XXXVI, 12.32 in CCHMC-DIPG-1, and 12.37 in SU-DIPG-XIII (Figure 3A-D). These findings demonstrate a synergistic effect of THIO and IR in suppressing DIPG cell proliferation. Notably, peak synergy was consistently observed at an IR dose of 4 Gy and THIO concentrations at or below the respective IC_50_ values for each cell line (Figure 3A-D). We next selected 0.5 μM THIO and 4 Gy IR as rational combination treatment doses for subsequent mechanistic analyses. In both SU-DIPG-XXXVI and CCHMC-DIPG-1 cells, combined THIO and IR treatment resulted in a significantly greater accumulation of cells in G_2_/M phase compared with either treatment alone (Figure 3E-F). Consistent with these findings, combination treatment also significantly increased DNA damage, as assessed by γH2AX staining, and apoptotic cell death, as measured by cleaved caspase-3 staining, in both DIPG cell lines (Figure 3G-N). Collectively, these findings support a model in which THIO-induced telomeric damage establishes a pre-existing state of genomic stress that sensitizes tumor cells to subsequent radiation-induced DNA lesions, resulting in an accumulation of genomic and telomeric damage that exceeds the adaptive capacity of DIPG cells and ultimately promotes sustained cell cycle arrest and apoptotic cell death.

**Figure 3:**
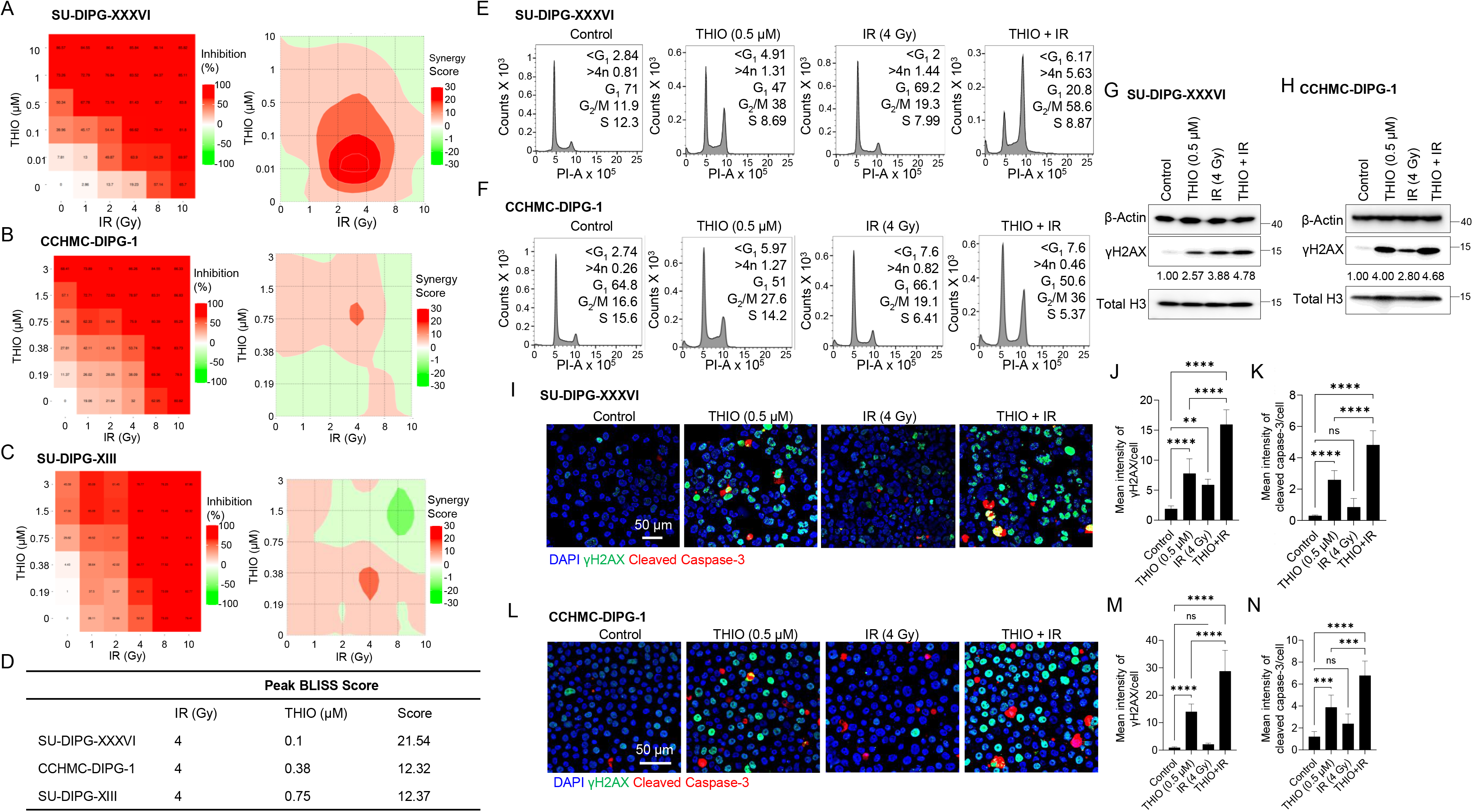
THIO synergizes with radiation to enhance DNA damage and promote apoptosis. **(A-C)** THIO-radiation combination inhibition matrices and corresponding Bliss Scoring landscapes in SU-DIPG-XXXVI, CCHMC-DIPG-1, and SU-DIPG-XIII cells. **(D)** Quantification of peak Bliss interaction scores across the indicated cell lines. **(E-F)** Cell cycle distribution assessed by flow cytometry following 48 hrs of treatment with vehicle, THIO (0.5 μM), IR (4 Gy), or the combination of THIO and IR. **(G-H)** Representative γH2AX immunoblot analysis following the indicated treatments. Densitometric values beneath the blots represent relative γH2AX protein levels normalized to total histone H3 and β-actin. **(I and L)** Representative immunofluorescence images showing DAPI (blue), γH2AX (green), and cleaved caspase-3 (red) in SU-DIPG-XXXVI, and CCHMC-DIPG-1 cells following 48 hrs of treatment with vehicle, THIO (0.5 μM), IR (4 Gy), or the combination of THIO and IR. **(J-K, M-N)** Quantification of γH2AX and cleaved caspase-3 fluorescence intensities using Image J. Mean fluorescence intensity was calculated from at least 3 fields per treatment condition and normalized to cell number, with 200-500 cells analyzed per treatment condition. All *in vitro* experiments were performed with n=2. Statistical significance is indicated as follows: **P < 0.01, ***P < 0.001, and ****P < 0.0001.

### THIO potentiates radiotherapeutic efficacy in orthotopic DIPG models

We next investigated whether the synergistic interaction between THIO and IR observed *in vitro* could be recapitulated *in vivo*. To this end, luciferase-expressing SU-DIPG-XXXVI cells were orthotopically implanted into the pons, and mice were randomized into treatment groups based on baseline BLI. The treatment regimen consisted of THIO administered at 5 mg/kg in combination of three cycles of fractionated RT (10 Gy per cycle, delivered as 2 Gy daily fractions), followed by continued intermittent THIO dosing until the experimental endpoint, as outlined in Figure 4A. Longitudinal BLI monitoring revealed progressive tumor growth in vehicle-treated mice, whereas THIO and IR monotherapies delayed tumor progression but failed to achieve sustained tumor control (Figure 4B, Figure S3A). Notably, combination treatment with THIO and IR resulted in significantly greater and more sustained suppression of tumor growth compared with either monotherapy (Figure 4B,). Consistent with these findings, combination-treated mice exhibited a pronounced survival benefit, with a median survival of 105.5 days compared with 37.5 days for vehicle-treated mice, 57 days for THIO-treated mice, and 47 days for IR-treated mice (Figure 4C). Furthermore, tumor tissue collected from mice at comparable time points demonstrated significantly increased cleaved caspase-3 staining following combination treatment relative to vehicle and either monotherapy, consistent with enhanced tumor cell apoptosis (Figure 4D-E). Of note, longitudinal body weight trajectories were comparable across treatment groups, supporting the tolerability of both THIO monotherapy and the THIO-IR combination regimen (Figure S3B). Collectively, these findings demonstrate that telomere-directed therapy substantially enhances the antitumor efficacy of fractionated RT and improves survival in an orthotopic DIPG model.

**Figure 4:**
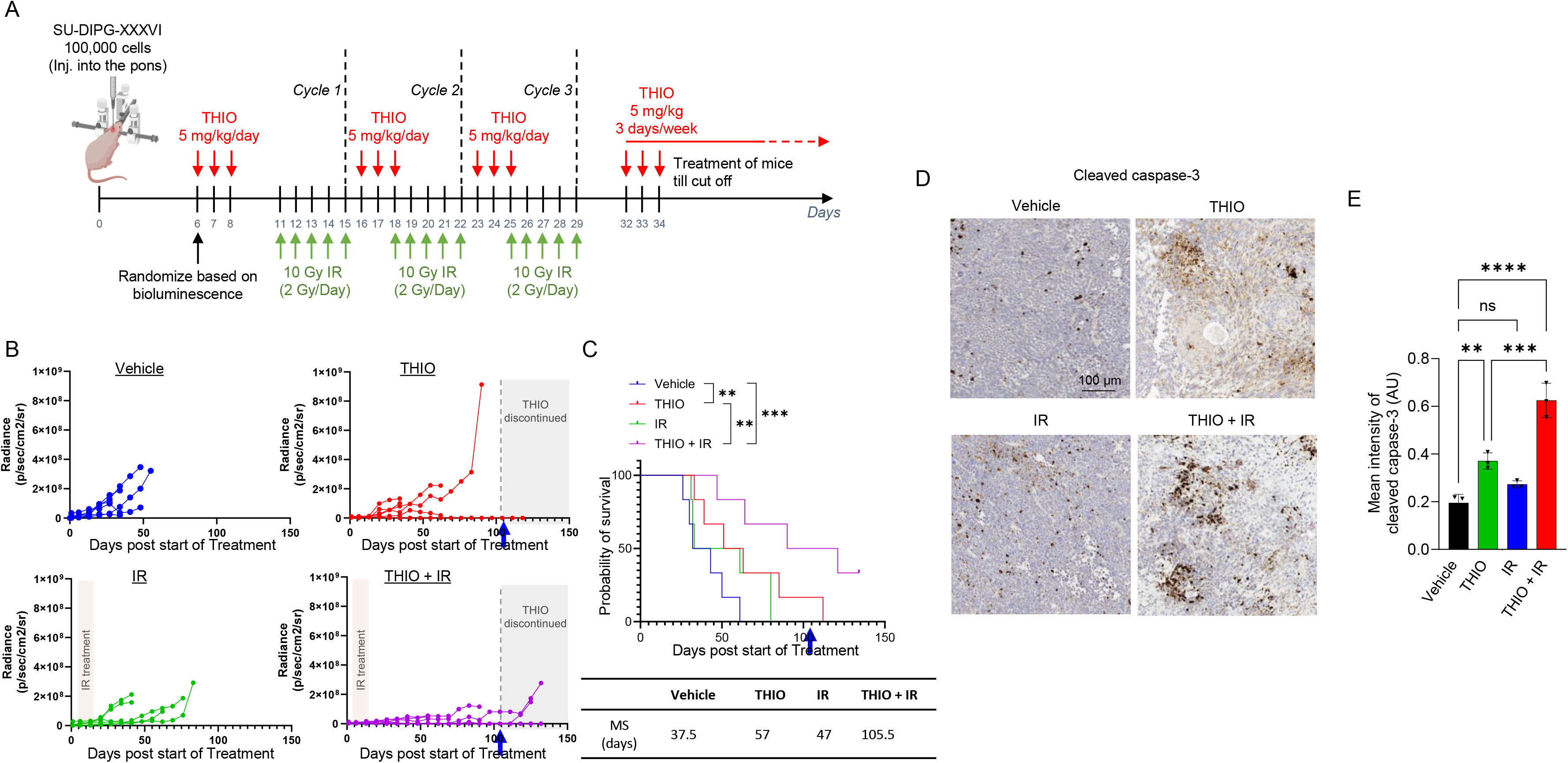
THIO enhances the therapeutic efficacy of fractionated radiation in orthotopic DIPG model. **(A)** Schematic representation of the orthotopic SU-DIPG-XXXVI treatment protocol. Mice received THIO (5 mg/kg) in combination with three cycles of fractionated radiation (10 Gy per cycle, 2 Gy/day, for a cumulative dose of 30 Gy), followed by intermittent THIO treatment. **(B)** Longitudinal monitoring of tumor burden by bioluminescence imaging using IVIS system. **(C)** Kaplan-Meier survival analysis of mice treated with vehicle, THIO, IR, or the combination of THIO and IR. Median survival was 37.5, 57, 47, and 105.5 days, respectively. . **(D)** Representative cleaved caspase-3 immunohistochemical staining and quantification of apoptosis in orthotopic tumor sections (n=3 mice per treatment group). Cleaved caspase-3 staining intensity was quantified using ImageJ and normalized to tumor area. Statistical significance is indicated as follows: **P < 0.01, ***P < 0.001, and ****P < 0.0001.

### THIO-induced telomere damage activates ATM/ATR signaling as an adaptive stress response

The sustained activation of ATM and ATR signaling following THIO treatment raises the possibility that these pathways function not only as markers of genomic and telomeric damage but also as adaptive survival mechanisms in DIPG cells. To investigate this possibility, we evaluated the effects of pharmacologic ATM or ATR inhibition in combination with THIO in SU-DIPG-XXXVI and CCHMC-DIPG-1 cells. The experimental timelines for early and delayed endpoint analyses are illustrated in Figure S4A. DIPG cells were pretreated with 10 µM ATM inhibitor (ATMi) or 50 nM ATR inhibitor (ATRi) for 2 hrs prior to exposure to 0.5 µM THIO (Figure S4A). Consistent with our previous findings (Figure 2E), THIO treatment alone induced ATM and ATR pathway activation, accompanied by increased γH2AX, although the relative contributions of these signaling pathways differed between the two DIPG cell lines (Figure 5A-B). Notably, in SU-DIPG-XXXVI cells, ATM inhibition was associated with increased ATR phosphorylation, whereas combined THIO and ATR inhibition resulted in enhanced CHK2 phosphorylation (Figure 5A). In CCHMC-DIPG-1 cells, combined THIO and ATR inhibition also resulted in a marked increase in γH2AX (Figure 5B). These findings suggest potential crosstalk between ATM and ATR signaling in response to genomic and telomeric damage, rather than a simple linear relationship between the two pathways. Furthermore, the attenuation of THIO-induced G_2_/M cell cycle arrest following either ATM or ATR inhibition suggests that these checkpoint kinases contribute to adaptive cellular responses that may promote survival following THIO treatment. Disruption of these checkpoint responses may allow cells harboring persistent genomic and telomeric lesions to progress through mitosis, thereby propagating unresolved DNA damage and potentially amplifying genomic instability, ultimately promoting apoptotic cell death (Figure 5C-D). Consistent with this model, the consequences of checkpoint kinase inhibition became more pronounced over time. Immunofluorescence analysis at early (day 2) and delayed (day 7) time points revealed limited apoptotic activity following checkpoint inhibitor monotherapy, whereas THIO-containing combination treatments resulted in significantly increased cleaved caspase-3 positivity by day 7 (Figure 5E-H). Additionally, γH2AX staining and quantification confirmed the persistence of DNA damage following these combination treatments (Figure 5E-F and S4B-C). Taken together, these findings support a model in which THIO treatment activates ATM/ATR-dependent checkpoint signaling, transiently restraining cell cycle progression and potentially enabling cellular adaptation to persistent genomic stress. Pharmacologic disruption of these pathways attenuates checkpoint arrest, potentially allowing damaged cells to progress through mitosis despite unresolved genomic and telomeric lesions. This may ultimately convert a sustained damage response into catastrophic genomic instability, culminating in mitotic catastrophe and enhanced apoptotic cell death. This mechanistic framework is consistent with the emerging therapeutic relevance of ATM and ATR dependencies in replication-stressed DMG^13,42,43^, and supports the concept that exacerbating replication-associated stress before introducing radiation-induced DNA damage may enhance therapeutic efficacy.

**Figure 5:**
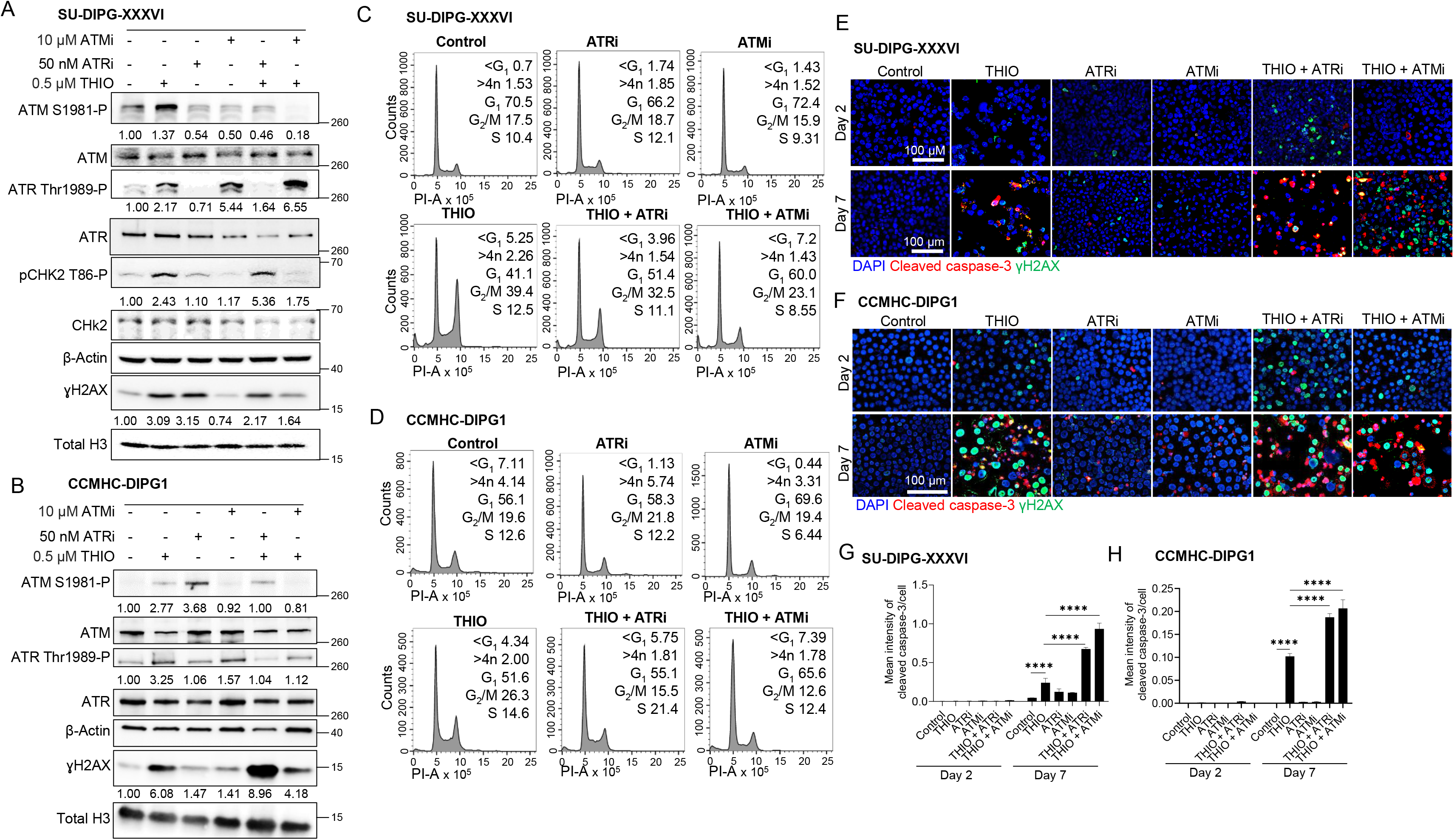
ATM and ATR checkpoint signaling modulates cellular response to THIO-induced telomere damage. **(A-B)** Immunoblot analysis of ATM, ATR, CHK2, and γH2AX in SU-DIPG-XXXVI and CCHMC-DIPG-1 cells following treatment with THIO (0.5 μM), an ATR inhibitor (50 nM), an ATM inhibitor (10 μM), or the indicated combination treatments. Densitometric values beneath the blots represent relative protein levels. Phosphorylated ATM, ATR, and CHK2 levels were normalized to their respective total protein levels and subsequently to β-actin. γH2AX levels were normalized to total histone H3. **(C-D)** Flow cytometric analysis of cell cycle distribution demonstrating the effects of ATM or ATR inhibition on THIO-induced cell cycle checkpoint arrest following treatment with THIO (0.5 μM), ATR inhibitor (50 nM), ATM inhibitor (10 μM), or the indicated combinations. **(E-F)** Representative immunofluorescence images showing DAPI (blue), γH2AX (green), and cleaved caspase-3 (red) staining at early and delayed time points in SU-DIPG-XXXVI and CCHMC-DIPG-1 cells following treatment with THIO (0.5 μM), ATR inhibitor (50 nM), ATM inhibitor (10 μM), or the indicated combinations. **(G-H)** Quantification of cleaved caspase-3 fluorescence intensity demonstrating enhanced delayed apoptosis following combined THIO and DDR pathway inhibition. Treatment schedules are detailed in Figure S4A. Cleaved caspase-3 fluorescence intensity was quantified using ImageJ, averaged across at least three fields per treatment condition, and normalized to cell number (200-500 cells analyzed per treatment condition). All *in vitro* experiments were performed with n=2. P values are indicated as follows: **P < 0.01, ***P < 0.001, and ****P < 0.0001.

## DISCUSSION

Our study identifies telomere integrity as a therapeutically exploitable determinant of radiation response in DIPG and provides a mechanistic rationale for combining telomere-directed therapy with radiotherapy. Our findings support a sequential model in which THIO induces telomerase-dependent telomeric damage, triggering progressive DDR signaling, suppression of proliferative transcriptional programs, and activation of cell cycle checkpoints. Subsequent radiation exposure in this pre-existing state of genomic and telomeric stress further amplifies DNA damage and promotes apoptotic cell death beyond the effects observed with either treatment alone. Importantly, pharmacologic inhibition of ATM or ATR signaling provides additional evidence that checkpoint activation in response to THIO-induced telomere damage may represent an adaptive survival mechanism that enables DIPG cells to temporarily restrain cell cycle progression and tolerate persistent genomic lesions, rather than simply serving as a marker of cellular injury. Thus, disrupting these compensatory checkpoint responses may further compromise the ability of DIPG cells to withstand telomere-directed damage and enhance their susceptibility to therapeutic interventions.

The mechanism of action of THIO fundamentally distinguishes it from conventional telomerase inhibitors, particularly Imetelstat (IMT). Direct telomerase inhibition relies on replication-dependent progressive telomere shortening, potentially requiring multiple rounds of cell division before telomeres reach a critically short length sufficient to trigger cellular senescence or apoptosis. This delayed onset of cytotoxicity may represent a significant therapeutic limitation in rapidly progressive pediatric brain tumors, where timely tumor control is essential. Indeed, our previous molecular studies and phase II clinical trials evaluating IMT as a direct telomerase inhibitor in children with CNS tumors highlighted limitations in its tolerability, with treatment-related toxicities restricting the feasibility of sustained or frequent dosing required to achieve progressive telomere attrition^26^. Similarly, in our previous preclinical study, IMT failed to produce durable radiosensitization in a high-risk medulloblastoma model despite inducing measurable telomere shortening^25^. In contrast, THIO exploits telomerase enzymatic activity to incorporate an abnormal guanosine analogue directly into newly synthesized telomeric DNA^27^. Biochemical studies have demonstrated that THIO incorporation impairs telomerase repeat-addition processivity^44^, while more recent work has shown that incorporation of THIO into telomeres promotes the formation of a nonproductive telomerase-telomere complex^45^. These mechanisms provide a potential explanation for the progressive accumulation of TIFs observed in our DIPG models, consistent with ongoing telomeric damage resulting from THIO incorporation and impaired telomere maintenance.

Our findings indicate that the cellular response to THIO extends beyond telomere dysfunction itself, encompassing broad transcriptional and metabolic reprogramming. Specifically, THIO treatment suppressed *E2F* target genes, G_2_/M checkpoint-associated genes, mitotic spindle components, MYC-associated programs, oxidative phosphorylation, glycolysis, and lipid metabolic pathways, while concurrently activating p53 signaling, unfolded protein response, and broader cellular stress and inflammatory programs. Notably, the apparent discordance between the downregulation of G_2_/M-associated transcriptional signatures and the accumulation of cells in G_2_/M phase observed by flow cytometry is biologically informative rather than contradictory. These assays capture distinct aspects of the cellular response. Flow cytometry identifies the cell cycle phase in which cells accumulate, whereas transcriptomic pathway analysis reflects the activity of gene expression programs associated with cell cycle progression. Thus, THIO-treated cells may accumulate at the G_2_/M checkpoint while simultaneously exhibiting suppression of the transcriptional machinery required for productive cell cycle progression and mitotic entry. Similar phenomena have been described in other experimental systems, in which checkpoint activation coincides with suppression of G_2_/M-associated transcriptional networks and an increased proportion of cells arrested in G_2_/M phase^46^. Together, these findings support a model in which THIO-induced genomic, and telomeric stress activates G_2_/M checkpoint signaling while suppressing the transcriptional programs necessary for cell cycle progression. Consequently, damaged cells accumulate at the G_2_/M boundary but exhibit a diminished capacity to execute the proliferative and mitotic programs required for continued cell division.

The ATM and ATR signaling findings provide a mechanistic link between telomere dysfunction and radiosensitization. Dysfunctional telomeres are recognized as sites of DNA damage and can engage both ATM- and ATR-dependent signaling pathways. Notably, we observed distinct temporal patterns of pathway activation, with ATR phosphorylation exhibiting an acute response that gradually diminished over time, whereas ATM activation persisted and increased over the course of THIO treatment. This sustained ATM response suggests that ATM signaling may represent a predominant component of the cellular response to THIO-induced telomere dysfunction, consistent with our previous findings^28^. We and others have previously demonstrated that telomere dysfunction enhances sensitivity to IR, in part by altering the kinetics of the DDR, particularly through ATM activation^47,48^. Building on these findings, our current study demonstrates that pharmacological inhibition of ATR or ATM attenuates THIO-induced G_2_/M accumulation, followed by a pronounced apoptotic response (Figure 5E-H). This temporal sequence suggests that checkpoint disruption allows cells harboring extensive DNA damage to progress into an unsustainable cell-cycle state, ultimately triggering cell death. These findings are conceptually consistent with the broader rationale for combining DDR inhibitors with radiation in brain tumors^13^. Emerging evidence further supports the therapeutic potential of ATM inhibition in DMG, particularly with the brain-penetrant ATM inhibitor AZD1390, which has demonstrated robust blood–brain barrier penetration and radiosensitizing activity in preclinical brain tumor models^49–51^. The ongoing clinical evaluation of AZD1390 in combination with radiotherapy (NCT06894979 and NCT03970447) further underscores the translational relevance of targeting ATM signaling. Future studies will focus on long-term survival analyses, combination-response matrices, and orthotopic validation using clinically relevant, brain-penetrant checkpoint inhibitors to establish whether these interactions are synergistic and therapeutically actionable.

The interaction between THIO with RT is particularly relevant given that RT remains the standard of care for DMG, yet its therapeutic benefit is typically transient^7,8^. Our findings suggest that THIO does not simply introduce an independent cytotoxic insult but instead alters the cellular context in which radiation-induced damage occurs. Specifically, THIO-treated cells exhibit pre-existing telomeric lesions, activated DDR signaling, and suppression of proliferative transcriptional programs. Consequently, radiation superimposes genome-wide DNA damage on cells already operating near their threshold for damage tolerance. The resulting increases in γH2AX, G_2_/M accumulation, and apoptosis are consistent with enhanced radiosensitivity. Importantly, the marked extension of median survival to 105.5 days in the combination treatment group (Figure 4C) provides *in vivo* support for this therapeutic interaction.

In summary, our findings establish telomere-directed damage as a promising strategy for altering the therapeutic vulnerability of DMG. THIO simultaneously induces telomeric damage, suppresses proliferative programs, and engages ATM/ATR-dependent checkpoint signaling. Radiation exploits this pre-existing cellular stress to amplify DNA damage and improve tumor control, whereas checkpoint inhibition reveals an additional adaptive dependency that can be therapeutically exploited. Collectively, these findings support a model in which telomere targeting serves not only as a direct cytotoxic strategy but also as a therapeutic priming approach that enhances the vulnerability of DMG cells to DNA-damaging treatments. This framework provides a rationale for further investigating telomere-directed therapy in combination with radiotherapy and DDR-targeted agents as a strategy to improve therapeutic outcomes in DMG.

### Limitation of Study

Although intracranial efficacy and intratumoral TIF formation provide functional evidence of brain-tumor exposure, formal pharmacokinetic measurements were not included in the supplied dataset. Quantification of THIO or active metabolites in plasma, normal brain, and tumor would strengthen translational conclusions regarding CNS penetration.

## Acknowledgements

This work was supported by MAIA Biotechnology, Inc. (Chicago, IL) to R.D., by CancerFree KIDS Pediatric Cancer Research Alliance to B. U. and R. D., The Pray Hope Believe Foundation to R.D., and Nationwide Children’s Hospital.

We sincerely thank the Small Animal Core (SAIC) at the Ohio State University as well as the Animal Resource, Flow Cytometry, Histopathology, Microscopy, and the Steve Cindy Rasmussen Institute for Genomic Medicine Cores and Microscopy Core at Nationwide Children’s Hospital.

We thank Dr. Michelle Monje (Stanford University) for kindly providing SU-DIPG-XXXVI, SU-DIPG-IV, SU-DIPG-XIII. Dr. Timothy N. Phoenix (Cincinnati Children’s Hospital) for the kind gift of IUE-24-C5, IUE-24-B1, and IUE-26-D6, and Dr. James Olson (Seattle Children’s Hospital) for kindly providing PBT-29FH.

## Author contributions

Conceptualization: B.U., R.D.; Methodology and Experimental Design: B.U., Z.W., D.K.M., S.S.K., R.D.; Data Acquisition and Analysis: B.U., Z.W., A.M.I., K.K., H-H. P.; Patient Identification and Consent: M.F.; Writing Original Draft: B.U.; Writing, Review and Editing: All authors; Funding Acquisition: B.U., R.D., S.G.; Supervision: R.D.

## Data availability

All supporting data generated for this manuscript are provided in the supplemental material file. Any other supporting data is available upon request from the corresponding author.

## SUPPLEMENTARY FIGURE LEGENDS

**Figure S1: THIO sensitivity, cell-cycle effects, and telomere dysfunction in additional DIPG models.**

**(A)** Evaluation of *hTERT* expression by RT-qPCR in patient-derived DIPG cell lines. Error bars represent the SD of two independent experiments; each performed in triplicates.

**(B)** Assessment of telomerase activity in patient-derived DIPG cell lines by telomeric repeat amplification protocol (TRAP) assay. HeLa cells and CHAPS buffer were used as positive and negative controls, respectively.

**(C)** THIO dose-response curves in IUE-derived murine H3-wild-type, H3.3K27M, and H3.1K27M cells. Cell proliferation was assessed 72 hrs after THIO treatment.

**(D)** THIO IC_50_ values (µM) across H3-wild-type, H3.3K27M, and H3.1K27M patient-derived DIPG and mouse cell lines.

**(E)** Cell cycle analysis of CCHMC-DIPG-1 cells by flow cytometry following treatment with 1 µM THIO for 24-72 hrs. Quantification of cell-cycle phase distribution is shown on the right.

**(F)** Representative telomere IF-FISH images and quantification of TIFs in CCHMC-DIPG-1 cells following THIO treatment. Arrows indicate representative TIFs. The number of TIFs per cell was quantified across 200-500 cells per treatment condition. Error bars represent the SD of at least three fields per treatment condition, with more than 100 cells analyzed per field. *In vitro* experiments were performed with n=3. P values are indicated as ****P < 0.0001.

**Figure S2: Cross-model transcriptomic response to THIO treatment.**

**(A)** Principal-component analysis (PCA) of vehicle- and THIO-treated CCHMC-DIPG-1 and SU-DIPG-XXXVI cells, illustrating global transcriptional differences across treatment conditions and cell models.

**(B)** Cross-model comparison of THIO-induced log_2_ fold changes in CCHMC-DIPG-1 and SU-DIPG-XXXVI cells, identifying genes that were concordantly upregulated, concordantly downregulated, or differentially regulated between the two models.

**Figure S3: Longitudinal tumor imaging and body-weight monitoring during *in vivo* THIO and radiation treatment.**

**(A)** Representative serial bioluminescence images acquired by IVIS of mice bearing orthotopic SU-DIPG-XXXVI tumors following treatment with vehicle, THIO, radiation, or the combination of THIO and radiation.

**(B)** Longitudinal changes in mean body weight across the four treatment groups. Shaded regions indicate the treatment and post-treatment periods.

**Figure S4: Experimental design and γH2AX response to THIO treatment in combination with ATM or ATR inhibition.**

**(A)** Schematic of the treatment schedule and experimental endpoints for SU-DIPG-XXXVI and CCHMC-DIPG-1 cells. Cells were pretreated with an ATM or ATR inhibitor for 2 hrs before THIO exposure. Analyses were performed on day 2 to assess early response and day 7 to evaluate delayed response.

**(B-C)** Quantification of mean γH2AX intensity per cell at day 2 and day 7 across control, THIO, ATR inhibitor, ATM inhibitor, THIO + ATR inhibitor, and THIO + ATM inhibitor conditions. γH2AX fluorescence intensity was quantified using ImageJ. Mean intensity values were calculated from at least three fields per treatment condition and normalized to the number of cells analyzed (200-500 cells per treatment condition). *In vitro* experiments were performed with n=2. P values are indicated as **P < 0.01, ***P < 0.001, and ****P < 0.0001.

## REFERENCES

1. Fisher PG, Breiter SN, Carson BS, et al. A clinicopathologic reappraisal of brain stem tumor classification. Identification of pilocystic astrocytoma and fibrillary astrocytoma as distinct entities. Cancer. 2000; 89(7):1569–1576.

2. Mackay A, Burford A, Carvalho D, et al. Integrated Molecular Meta-Analysis of 1,000 Pediatric High-Grade and Diffuse Intrinsic Pontine Glioma. Cancer Cell. 2017; 32(4):520–537 e525.

3. Patil N, Kelly ME, Yeboa DN, et al. Epidemiology of brainstem high-grade gliomas in children and adolescents in the United States, 2000-2017. Neuro Oncol. 2021; 23(6):990–998.

4. Hoffman LM, Veldhuijzen van Zanten SEM, Colditz N, et al. Clinical, Radiologic, Pathologic, and Molecular Characteristics of Long-Term Survivors of Diffuse Intrinsic Pontine Glioma (DIPG): A Collaborative Report From the International and European Society for Pediatric Oncology DIPG Registries. J Clin Oncol. 2018; 36(19):1963–1972.

5. Caretti V, Bugiani M, Freret M, et al. Subventricular spread of diffuse intrinsic pontine glioma. Acta Neuropathol. 2014; 128(4):605–607.

6. Qin EY, Cooper DD, Abbott KL, et al. Neural Precursor-Derived Pleiotrophin Mediates Subventricular Zone Invasion by Glioma. Cell. 2017; 170(5):845–859 e819.

7. Haas-Kogan D, Indelicato D, Paganetti H, et al. National Cancer Institute Workshop on Proton Therapy for Children: Considerations Regarding Brainstem Injury. Int J Radiat Oncol Biol Phys. 2018; 101(1):152–168.

8. Hayashi A, Ito E, Omura M, et al. Hypofractionated radiotherapy in children with diffuse intrinsic pontine glioma. Pediatr Int. 2020; 62(1):47–51.

9. Arrillaga-Romany I, Gardner SL, Odia Y, et al. ONC201 (Dordaviprone) in Recurrent H3 K27M-Mutant Diffuse Midline Glioma. J Clin Oncol. 2024; 42(13):1542–1552.

10. Rechberger JS, Lu VM, Zhang L, Power EA, Daniels DJ. Clinical trials for diffuse intrinsic pontine glioma: the current state of affairs. Childs Nerv Syst. 2020; 36(1):39–46.

11. Vitanza NA, Monje M. Diffuse Intrinsic Pontine Glioma: From Diagnosis to Next-Generation Clinical Trials. Curr Treat Options Neurol. 2019; 21(8):37.

12. Hargrave D, Bartels U, Bouffet E. Diffuse brainstem glioma in children: critical review of clinical trials. Lancet Oncol. 2006; 7(3):241–248.

13. Rahman R, Shi DD, Reitman ZJ, et al. DNA damage response in brain tumors: A Society for Neuro-Oncology consensus review on mechanisms and translational efforts in neuro-oncology. Neuro Oncol. 2024; 26(8):1367–1387.

14. Pedersen H, Schmiegelow K, Hamerlik P. Radio-Resistance and DNA Repair in Pediatric Diffuse Midline Gliomas. Cancers (Basel). 2020; 12(10).

15. Blackburn EH. Structure and function of telomeres. Nature. 1991; 350(6319):569–573.

16. Bakkenist CJ, Drissi R, Wu J, Kastan MB, Dome JS. Disappearance of the telomere dysfunction-induced stress response in fully senescent cells. Cancer Res. 2004; 64(11):3748–3752.

17. d’Adda di Fagagna F, Reaper PM, Clay-Farrace L, et al. A DNA damage checkpoint response in telomere-initiated senescence. Nature. 2003; 426(6963):194–198.

18. Takai H, Smogorzewska A, de Lange T. DNA damage foci at dysfunctional telomeres. Curr Biol. 2003; 13(17):1549–1556.

19. Kim NW, Piatyszek MA, Prowse KR, et al. Specific association of human telomerase activity with immortal cells and cancer. Science. 1994; 266(5193):2011–2015.

20. Shay JW, Bacchetti S. A survey of telomerase activity in human cancer. Eur J Cancer. 1997; 33(5):787–791.

21. Dorris K, Sobo M, Onar-Thomas A, et al. Prognostic significance of telomere maintenance mechanisms in pediatric high-grade gliomas. J Neurooncol. 2014; 117(1):67–76.

22. Lulla RR, Buxton A, Krailo MD, et al. Vorinostat, temozolomide or bevacizumab with irradiation and maintenance BEV/TMZ in pediatric high-grade glioma: A Children’s Oncology Group Study. Neurooncol Adv. 2024; 6(1):vdae035.

23. Dikmen ZG, Gellert GC, Jackson S, et al. In vivo inhibition of lung cancer by GRN163L: a novel human telomerase inhibitor. Cancer Res. 2005; 65(17):7866–7873.

24. Hochreiter AE, Xiao H, Goldblatt EM, et al. Telomerase template antagonist GRN163L disrupts telomere maintenance, tumor growth, and metastasis of breast cancer. Clin Cancer Res. 2006; 12(10):3184–3192.

25. Sengupta S, Senthil Kumar S, Bondra K, Sobo M, Mo X, Drissi R. Limitations of radiosensitization by direct telomerase inhibition to treat high-risk medulloblastoma. Front Oncol. 2023; 13:1104670.

26. Salloum R, Hummel TR, Kumar SS, et al. A molecular biology and phase II study of imetelstat (GRN163L) in children with recurrent or refractory central nervous system malignancies: a pediatric brain tumor consortium study. J Neurooncol. 2016; 129(3):443–451.

27. Mender I, Gryaznov S, Dikmen ZG, Wright WE, Shay JW. Induction of telomere dysfunction mediated by the telomerase substrate precursor 6-thio-2’-deoxyguanosine. Cancer Discov. 2015; 5(1):82–95.

28. Sengupta S, Sobo M, Lee K, et al. Induced Telomere Damage to Treat Telomerase Expressing Therapy-Resistant Pediatric Brain Tumors. Mol Cancer Ther. 2018; 17(7):1504–1514.

29. Eglenen-Polat B, Kowash RR, Huang HC, et al. A telomere-targeting drug depletes cancer initiating cells and promotes anti-tumor immunity in small cell lung cancer. Nat Commun. 2024; 15(1):672.

30. Mender I, Siteni S, Barron S, et al. Activating an Adaptive Immune Response with a Telomerase-Mediated Telomere Targeting Therapeutic in Hepatocellular Carcinoma. Mol Cancer Ther. 2023; 22(6):737–750.

31. Mender I, Zhang A, Ren Z, et al. Telomere Stress Potentiates STING-Dependent Anti-tumor Immunity. Cancer Cell. 2020; 38(3):400–411 e406.

32. Senthil Kumar S, Sengupta S, Zhu X, et al. Diffuse Intrinsic Pontine Glioma Cells Are Vulnerable to Mitotic Abnormalities Associated with BMI-1 Modulation. Mol Cancer Res. 2020; 18(11):1711–1723.

33. Kumar SS, Sengupta S, Lee K, et al. BMI-1 is a potential therapeutic target in diffuse intrinsic pontine glioma. Oncotarget. 2017; 8(38):62962–62975.

34. du Chatinier A, Meel MH, Das AI, et al. Generation of immunocompetent syngeneic allograft mouse models for pediatric diffuse midline glioma. Neurooncol Adv. 2022; 4(1):vdac079.

35. Patel SK, Hartley RM, Wei X, et al. Generation of diffuse intrinsic pontine glioma mouse models by brainstem-targeted in utero electroporation. Neuro Oncol. 2020; 22(3):381–392.

36. Umaru B, Mishra DK, Kumar SS, et al. BMI-1 modulation and trafficking during M phase in diffuse intrinsic pontine glioma. FEBS Open bio. 2026; 16(6):1213–1221.

37. Umaru B, Sengupta S, Senthil Kumar S, Drissi R. Alternative Lengthening of Telomeres in Pediatric High-Grade Glioma and Therapeutic Implications. Cancers (Basel). 2023; 15(12).

38. Wang Z, Kumar A, Umaru B, et al. Integrated multi-omics identifies lineage-dependent myeloid cells recruitment and the APP-CD74 axis as an immunoregulatory target in pediatric high-grade glioma. Acta Neuropathol. 2026; 152(1).

39. Zhu X, Lazow MA, Schafer A, et al. A pilot radiogenomic study of DIPG reveals distinct subgroups with unique clinical trajectories and therapeutic targets. Acta Neuropathol Commun. 2021; 9(1):14.

40. Love MI, Huber W, Anders S. Moderated estimation of fold change and dispersion for RNA-seq data with DESeq2. Genome Biol. 2014; 15(12):550.

41. Ianevski A, Giri AK, Aittokallio T. SynergyFinder 3.0: an interactive analysis and consensus interpretation of multi-drug synergies across multiple samples. Nucleic Acids Res. 2022; 50(W1):W739–W743.

42. Hains AE, Chetal K, Nakatani T, et al. Multi-omics approaches reveal that diffuse midline gliomas present altered DNA replication and are susceptible to replication stress therapy. Genome Biol. 2024; 25(1):319.

43. Pal S, Kaplan JP, Nguyen H, et al. A druggable addiction to de novo pyrimidine biosynthesis in diffuse midline glioma. Cancer Cell. 2022; 40(9):957–972 e910.

44. Sanford SL, Welfer GA, Freudenthal BD, Opresko PL. Mechanisms of telomerase inhibition by oxidized and therapeutic dNTPs. Nat Commun. 2020; 11(1):5288.

45. Sanford SL, Badstubner M, Gerber M, et al. Therapeutic 6-thio-deoxyguanosine inhibits telomere elongation in cancer cells by inducing a non-productive stalled telomerase complex. Nat Commun. 2025; 17(1):6.

46. Tyagi AK, Singh RP, Agarwal C, Chan DC, Agarwal R. Silibinin strongly synergizes human prostate carcinoma DU145 cells to doxorubicin-induced growth Inhibition, G2-M arrest, and apoptosis. Clin Cancer Res. 2002; 8(11):3512–3519.

47. Drissi R, Wu J, Hu Y, Bockhold C, Dome JS. Telomere shortening alters the kinetics of the DNA damage response after ionizing radiation in human cells. Cancer Prev Res (Phila). 2011; 4(12):1973–1981.

48. Wong KK, Chang S, Weiler SR, et al. Telomere dysfunction impairs DNA repair and enhances sensitivity to ionizing radiation. Nat Genet. 2000; 26(1):85–88.

49. Durant ST, Zheng L, Wang Y, et al. The brain-penetrant clinical ATM inhibitor AZD1390 radiosensitizes and improves survival of preclinical brain tumor models. Sci Adv. 2018; 4(6):eaat1719.

50. GBM AGILE: Global Adaptive Trial Master Protocol: An International, Seamless Phase II/III Response Adaptive Randomization Platform Trial Designed To Evaluate Multiple Regimens In Newly Diagnosed and Recurrent GBM. 2019. https://clinicaltrials.gov/study/NCT03970447.

51. A Phase I Study to Assess the Safety, Tolerability, Pharmacokinetics, and Preliminary Efficacy of AZD1390 (NSC# 852149) When Combined With Focal Radiation in Pediatric Patients With High Grade Glioma. 2025. https://clinicaltrials.gov/study/NCT06894979.

