## Supplementary material for "Targeting Telomere Integrity Enhances Radiosensitivity and Reveals Adaptive DNA Damage Response Vulnerabilities in Diffuse Midline Glioma": Figures S1-S4

**Fig. S1**

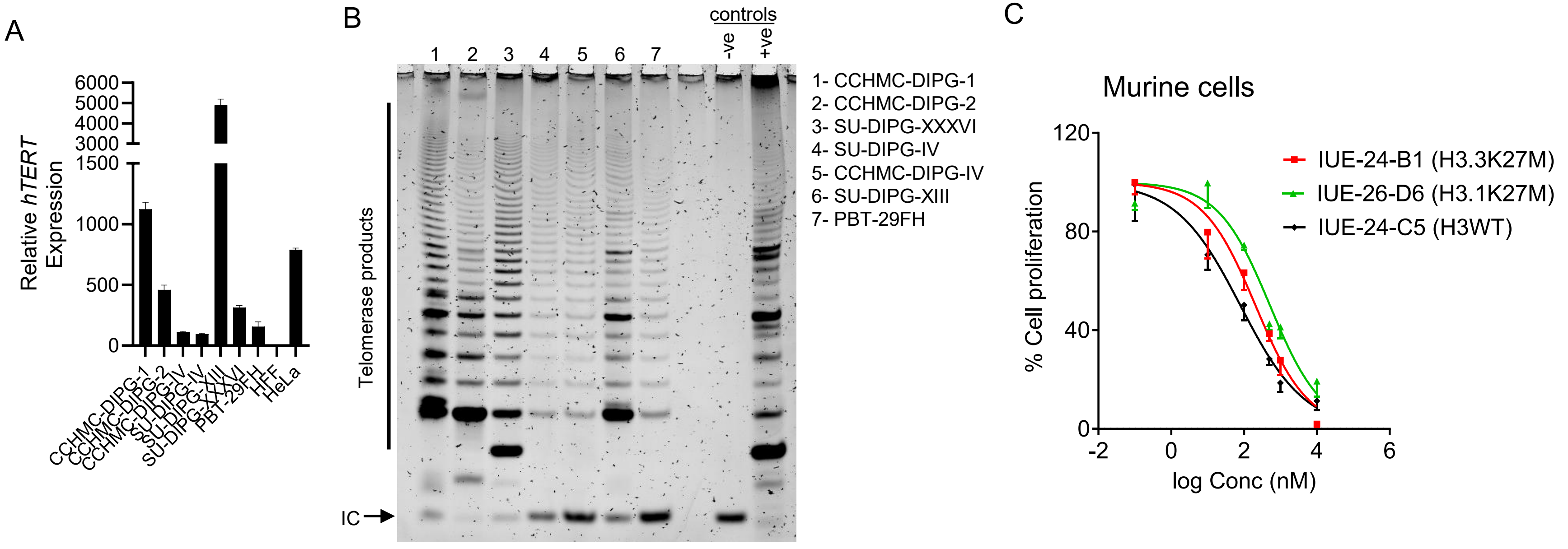


D

| Cell Line | $IC_{50}$ (µM) |
|---|---|
| CCHMC-DIPG-1 | 0.548 |
| CCHMC-DIPG-2 | 0.417 |
| CCHMC-DIPG-IV | 0.221 |
| SU-DIPG-IV | 0.597 |
| SU-DIPG-XIII | 0.996 |
| SU-DIPG-XXXVI | 0.198 |
| PBT-29FH | 0.135 |
| IUE-24-C5 | 0.080 |
| IUE-24-B1 | 0.208 |
| IUE-26-D6 | 0.518 |
| HFF | 14.79 |

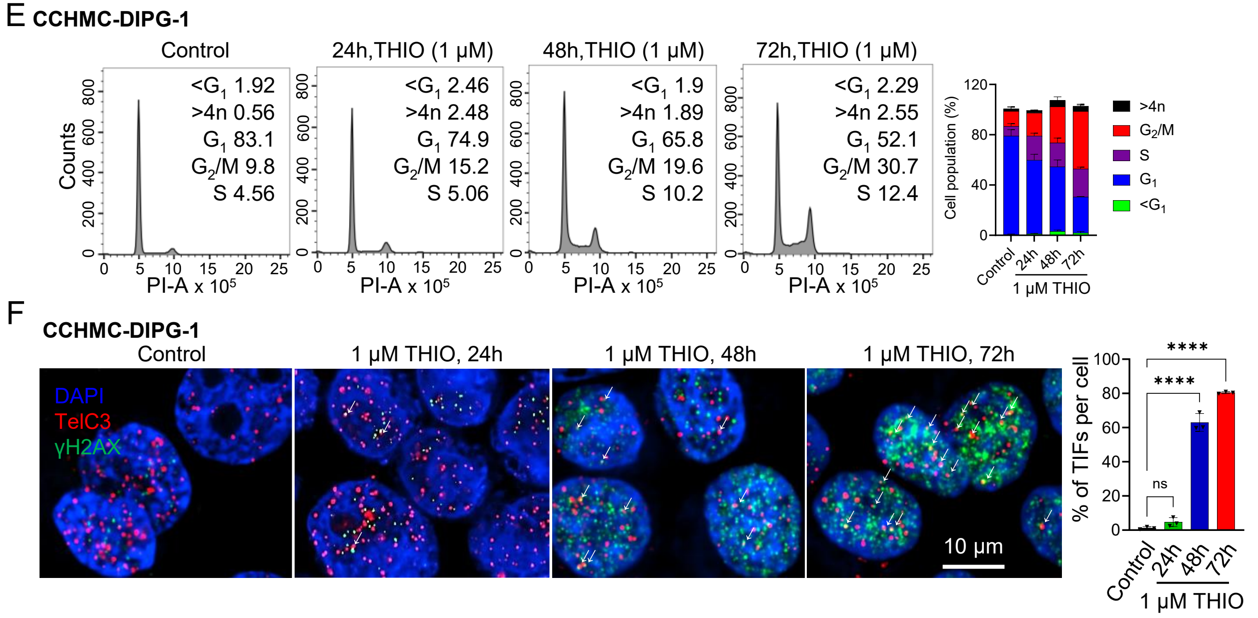

Fig. S2

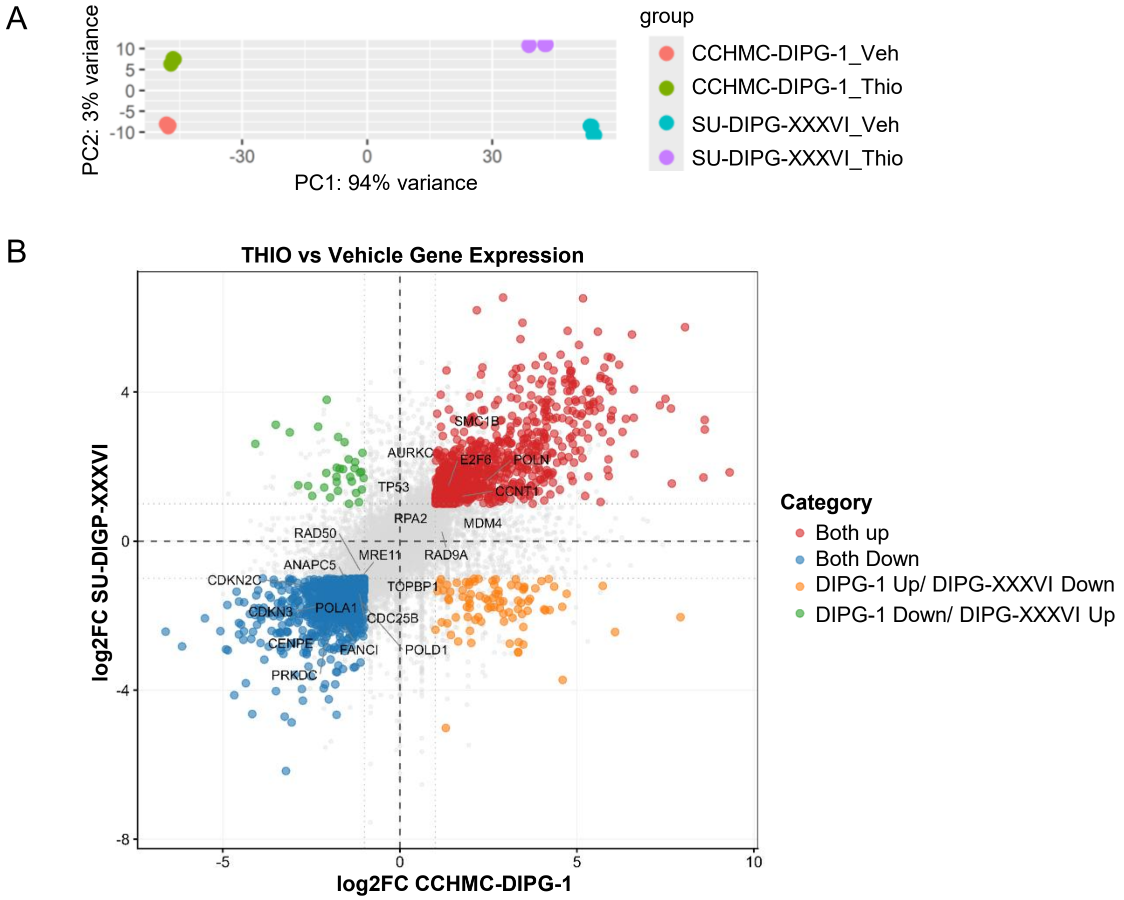
A
PC2: 3% variance
PC1: 94% variance
group
CCHMC-DIPG-1_Veh
CCHMC-DIPG-1_Thio
SU-DIPG-XXXVI_Veh
SU-DIPG-XXXVI_Thio
B
THIO vs Vehicle Gene Expression
log2FC SU-DIGP-XXXVI
log2FC CCHMC-DIPG-1
SMC1B
AURKC
E2F6
POLN
TP53
CCNT1
RPA2
MDM4
RAD50
MRE11
RAD9A
CDKN2C
ANAPC5
TOPBP1
CDKN3
POLA1
CDC25B
CENPE
FANCI
POLD1
PRKDC
Category
Both up
Both Down
DIPG-1 Up/ DIPG-XXXVI Down
DIPG-1 Down/ DIPG-XXXVI Up

### Fig. S3

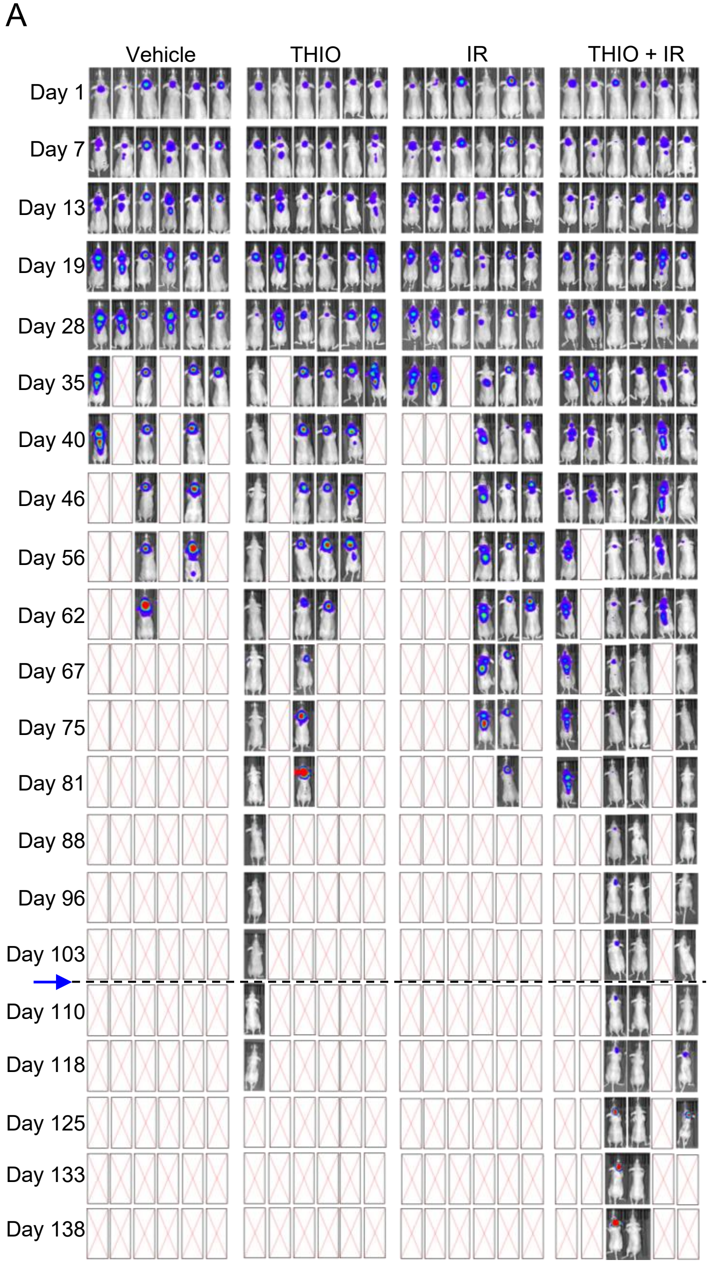
A
Vehicle
THIO
IR
THIO + IR
Day 1
Day 7
Day 13
Day 19
Day 28
Day 35
Day 40
Day 46
Day 56
Day 62
Day 67
Day 75
Day 81
Day 88
Day 96
Day 103
Day 110
Day 118
Day 125
Day 133
Day 138

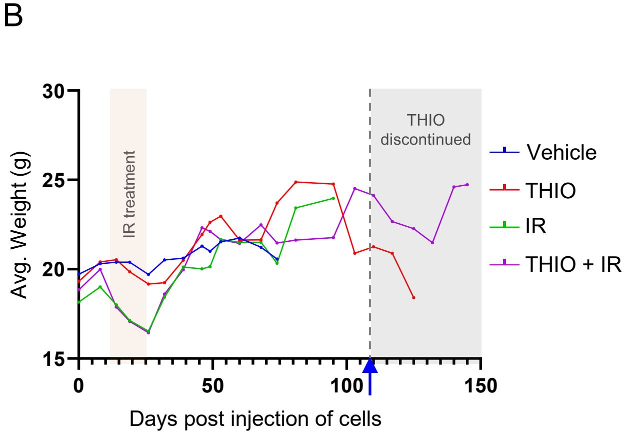
B
Avg. Weight (g)
15
20
25
30
IR treatment
THIO discontinued
0
50
100
150
Days post injection of cells
Vehicle
THIO
IR
THIO + IR

#### Fig. S4

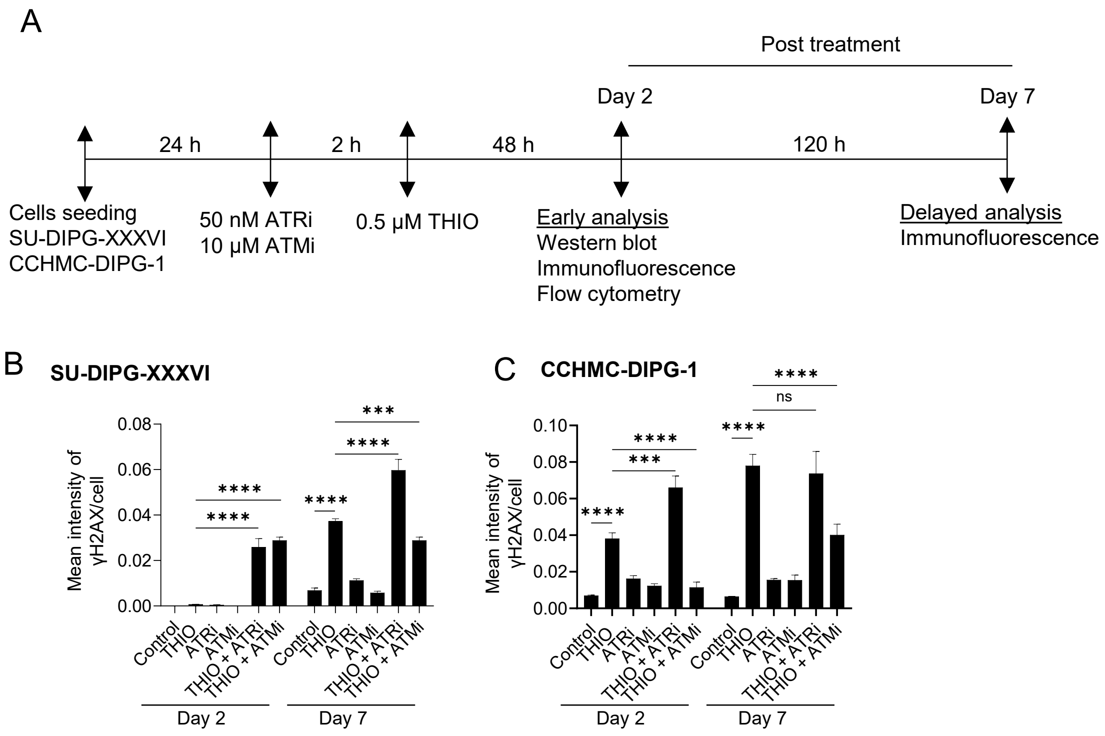

A
Post treatment
Day 2
Day 7
24 h
2 h
48 h
120 h
Cells seeding
SU-DIPG-XXXVI
CCHMC-DIPG-1
50 nM ATRi
10 μM ATMi
0.5 μM THIO
Early analysis
Western blot
Immunofluorescence
Flow cytometry
Delayed analysis
Immunofluorescence
B
SU-DIPG-XXXVI
Mean intensity of
γH2AX/cell
0.08
0.06
0.04
0.02
0.00
****
***
Control
THIO
ATRi
ATMi
THIO + ATRi
THIO + ATMi
Day 2
Day 7
C
CCHMC-DIPG-1
0.10
0.08
0.06
0.04
0.02
0.00
****
***
ns
Mean intensity of
γH2AX/cell
Control
THIO
ATRi
ATMi
THIO + ATRi
THIO + ATMi
Day 2
Day 7
